# A Novel Swine Model of the Acute Respiratory Distress Syndrome Using Clinically-Relevant Injury Exposures

**DOI:** 10.1101/2021.01.24.427964

**Authors:** Mohamad Hakam Tiba, Brendan M. McCracken, Danielle C. Leander, Carmen I. Colmenero, Jean A. Nemzek, Michael W. Sjoding, Kristine E. Konopka, Thomas L. Flott, J. Scott VanEpps, Rodney Daniels, Kevin R. Ward, Kathleen A. Stringer, Robert P. Dickson

## Abstract

To date, existing animal models of the acute respiratory distress syndrome (ARDS) have failed to translate preclinical discoveries into effective pharmacotherapy or diagnostic biomarkers. To address this translational gap, we developed a high-fidelity swine model of ARDS utilizing clinically-relevant lung injury exposures. Fourteen male swine were anesthetized, mechanically ventilated, and surgically instrumented for hemodynamic monitoring, blood, and tissue sampling. Animals were allocated to one of three groups: 1) *Indirect lung injury only*: animals were inoculated by direct injection of *E. coli* into the kidney parenchyma, provoking systemic inflammation and distributive shock physiology; 2) *Direct lung injury only*: animals received volutrauma, hyperoxia, and bronchoscope-delivered gastric particles; 3) *Combined indirect and direct lung injury:* animals were administered both above-described indirect and direct lung injury exposures. Animals were monitored for up to 12 hours, with serial collection of physiologic data, blood samples, and radiographic imaging. Lung tissue was acquired post-mortem for pathological examination. In contrast to *indirect lung injury only* and *direct lung injury only* groups, animals in the *combined indirect and direct lung injury* group exhibited all of the physiological, radiographic, and histopathologic hallmarks of human ARDS: impaired gas exchange (mean PaO_2_/FiO_2_ ratio 124.8 ± 63.8), diffuse bilateral opacities on chest radiographs, and extensive pathologic evidence of diffuse alveolar damage. Our novel porcine model of ARDS, built on clinically-relevant lung injury exposures, faithfully recapitulates the physiologic, radiographic, and histopathologic features of human ARDS, and fills a crucial gap in the translational study of human lung injury.

## Introduction

The acute respiratory distress syndrome (ARDS) is a life-threatening lung condition that affects more than 200,000 people in the United States each year with a mortality rate of approximately 40%(1, 2). As a clinically-defined syndrome, ARDS has undergone only modest refinement in its definition since its first report in 1967(3–5). Despite a half century of experimental and clinical study, the diagnosis of ARDS remains entirely clinical (with no molecular biomarkers), and its management remains entirely supportive (with no targeted therapies).

A major barrier to advances in our diagnosis and management of ARDS has been reliance on inadequate preclinical animal models to study the syndrome(6). The vast majority of experimental research on ARDS has been performed using small animal (i.e., rodent) models(7). This reliance on rodent modeling of ARDS has not been due to their fidelity to human disease, but rather due to ease of handling, cost, accessible reagents, and availability of purebred and genetically engineered strains. Anatomically, murine lungs have a distinct lobar structure with far fewer branching airways than humans, extensive bronchial-associated lymphoid tissue, and a near-absence of submucosal glands(7). Mice also profoundly differ from humans in their innate and adaptive immune response to injury, including fewer circulating neutrophils, absence of defensins, and a distinct chemokine repertoire(8). Notably, murine lungs almost never form hyaline membranes, a histopathological hallmark of diffuse alveolar damage (the histopathological hallmark of human ARDS)(9). For these reasons, the 2011 American Thoracic Society workshop report on experimental acute lung injury conceded “the responses of animal [murine] and human lungs to an injurious stimulus cannot be expected to be identical or perhaps even similar.”(7) Additionally, rodent models almost all preclude the study of co-interventions and organ support (e.g. vasopressors or mechanical ventilation), serial sampling across anatomic compartments, or radiographic study. For these reasons, the NHLBI has identified the need for large-animal models of ARDS as a research priority(10).

In contrast to rodent models, swine models of ARDS represent a promising alternative. The swine lung exhibits lobar, interlobular, and airway anatomy similar to that of humans(11), and immune gene expression of swine is far more similar to that of humans(12–17). The metabolite composition of swine lung tissue is far more representative of human lungs than is that of rodent species(15). Several swine models of ARDS exist, yet these rely on clinically unrepresentative single exposures (e.g. oleic acid infusion(18, 19), surfactant washout(20)). To our knowledge, no existing swine model recapitulates the core features of human ARDS using clinically-relevant exposures.

To address these gaps, we sought to establish a preclinical model of ARDS using clinically-relevant exposures that 1) faithfully recapitulates the physiologic, radiographic, and histopathologic features of human ARDS, 2) allows for longitudinal study of pathogenesis, underlying mechanisms, and treatment strategies, and 3) permits study of co-interventions and organ support (e.g. vasopressors and mechanical ventilation). Motivated by clinical(21) and experimental(22) observations that both epithelial *and* endothelial injury are necessary to provoke the full pathophysiologic and clinical manifestations of ARDS, we hypothesized that a combination of indirect lung injury (sepsis(23)) and direct lung injury (concurrent administration of volutrauma, hyperoxia, and instillation of gastric particles into the airways) would be required to induce all of the clinical and biological hallmarks of human ARDS. Selective data from the *indirect lung injury only* (sepsis) group has previously been published(23).

## Methods

This study adhered to the principles stated in the *Guide for the Care and Use of Laboratory Animals* (24), and was approved by the Institutional Animal Care and Use Committee (IACUC). Animals were acquired through an IACUC-approved vendor and acclimated for 5-10 days before experimentation.

### Anesthesia and Surgical Instrumentation

Fourteen male Yorkshire-mix swine, approximately 14-16 weeks of age, were fasted overnight with *ad libitum* access to water. Anesthesia was induced using an intramuscular injection of ketamine/xylazine combination (22mg/kg & 2mg/kg) followed by 1.5-2.5 % isoflurane administered through a facemask. Animals were intubated using a 7.5 mm cuffed endotracheal tube and mechanically ventilated to maintain end-tidal CO_2_ (PetCO_2_) at 35-45 mmHg. Heart rate (HR), electrocardiograph (ECG), PetCO_2_ and pulse oximeter oxygen saturation (SpO_2_) were monitored using a veterinary monitor (Surgivet advisor, Smiths Medical, St. Paul, MN). Body temperature was maintained between 37-38.5°C using a feedback-controlled warming blanket (Cincinnati SubZero, Blanketrol II, Cincinnati, OH).

Under aseptic conditions, the right carotid artery, the right external and internal jugular veins were cannulated to provide continuous monitoring of mean arterial blood pressure (MAP), pulmonary artery pressure (PAP), heart rate, core temperature, arterial and mixed venous blood sampling as well as for intravenous anesthetic administration. A midline laparotomy was performed to access the bladder, the left kidney, isolate the ureter, and for placement of an indwelling Foley catheter for urine draining. At the end of surgical instrumentation, inhalant anesthesia was transitioned to total intravenous anesthesia by continuous infusions of midazolam (5-20 mcg/kg/min), fentanyl (0.03-4.0 mcg/kg/min), and propofol (10-100 mcg/kg/min) for the remainder of the experiment. Baseline hemodynamic metrics and blood samples for hematology, serum chemistry, and arterial and venous lactate, glucose, electrolytes, oximetry and blood gasses were obtained (**Table 1**). Ventral-dorsal thoracic radiographic images were obtained using a veterinary portable X-ray (minxray hf100+, minxray, Northbrook, IL). A 5 mL inoculum (*E. coli* culture or saline sham) was administered into the left kidney parenchyma over 15 minutes (0.33mL/min) and post-injection procedures were done as previously described(23). Completion of the renal inoculation was considered Time 0 (T_0_). The abdominal wall and skin were closed in layers. The ureter was occluded for a duration of 1 hour and then unoccluded. Intravenous crystalloids were administered starting at T_2_ (7.5-10 ml/kg/hr) and continued for the duration of experimentation.

**Table 1.** Baseline characteristics by group. Data are presented as mean (standard deviation). Statistical significance was set at α < 0.05. * Denotes statistically significant difference between Group 1 and Group 2. ^ Denotes statistically significant difference between Group 1 and Group 3. PaO_2_, Partial pressure of oxygen. FiO_2_, Fraction of inspired oxygen. SmvO_2_, Mixed venous oxygen saturation (%); PetCO_2_, End-tidal CO_2_ (mmHg). SaO2, Arterial oxygen saturation. PaCO_2_, Partial pressure of arterial CO_2_.

| Characteristic | Experimental Group |  |  |
| --- | --- | --- | --- |
|  | Group 1:<br>*Indirect lung injury only* | Group 2:<br>*Direct lung injury only* | Group 3:<br>*Combined indirect and direct lung injury* |
| n | 5 | 4 | 5 |
| Weight (kg) | 43(3.81) | 44(2.06) | 43(0.71) |
| Mean Arterial Pressure (mmHg) | 85.5(9.31) | 90.1(15.13) | 93.0(7.19) |
| Pulmonary Artery Pressure (mmHg) | 21.3(7.38) | 13.5(6.23) | 18.6(2.90) |
| Heart Rate (BPM) | 74(5.8) | 78(6.5) | 70(9.3) |
| Temperature(°C) | 37.8(0.80) | 37.2(0.84) | 37.4(0.61) |
| pH | 7.474(0.022) | 7.401(0.081) | 7.442(0.093) |
| Lactate (mEq/L) | 1.4(0.95) | 0.5(0.05) | 0.7(0.36) |
| SaO<sub>2</sub> (%) | 98.2(1.28) | 100(0.00) | 100(0.00) |
| SmvO<sub>2</sub> (%) | 58.8(1.92)*^ | 75.2(6.26) | 78.8(4.62) |
| PaO<sub>2</sub>/FiO<sub>2</sub> Ratio | 470(47.9) | 454(28.8) | 494(67.4) |
| PetCO<sub>2</sub> (mmHg) | 43.9(3.92) | 39.9(2.15) | 39.0(3.69) |
| PaCO<sub>2</sub> (mmHg) | 40.4(5.01) | 46.3(6.76) | 43.7(6.68) |
| White Blood Count (10<sup>9</sup>/L) | 17.06(3.62) | 17.65(5.41) | 14.67(2.16) |
| Monocytes (10<sup>9</sup>/L) | 0.16(0.104) | 0.11(0.063) | 0.09(0.039) |
| Lymphocytes (10<sup>9</sup>/L) | 10.41(2.420) | 10.98(2.354) | 8.57(1.210) |
| Neutrophils (10<sup>9</sup>/L) | 6.48(2.270) | 6.56(3.363) | 6.00(1.276) |
| Blood Urea Nitrogen (mg/dL) | 7.4(3.43) | 5.5(1.29) | 7.6(2.40) |
| Creatinine (mg/dL) | 1.3(0.11)* | 1.0(0.14) | 1.2(0.19) |
| Hematocrit (%) | 40.6(6.55) | 38.1(2.83) | 35.6(3.07) |
| Platelet (10<sup>9</sup>/L) | 279(102.3) | 294(44.0) | 283(97.6) |

To ensure humane experimentation, our protocol included pre-specified criteria for experiment termination: 1) persistent low mean arterial pressure (<40 mmHg) for more than 2 hours, 2) persistent low PetCO_2_ (<25 mmHg) for more than 2 hours, 3) critical low mean arterial pressure (<25 mmHg) combined with critical low PetCO_2_ (<20mmHg), 4) critical low PaO_2_ (<55 mmHg) for more than 1 hour, 5) ventricular fibrillation/tachycardia, and 6) malignant hyperthermia due to inhalant anesthetics.

### Experimental Groups and Exposures

Animals were designated into one of three experimental groups as follows:

#### Group 1 – *Indirect lung injury only*

As recently described(23), a total volume of 5 ml containing an average culture count of 2.5×10^11^ CFUs of live *E. coli* was administered into the kidney’s parenchyma. No antibiotics were administered. Tidal volume (Vt) was set between 7-8 ml/kg using 21% fractional inspired O_2_ (FiO_2_) and 5 cmH_2_O of positive end-expiratory pressure (PEEP). As previously reported(23), this exposure provokes systemic inflammation and distributive shock physiology characteristic of sepsis.

#### Group 2- *Direct lung injury only*

1) <u>Volutrauma</u>: Tidal volume was set between 12-15 ml/kg during instrumentation and continued for the duration of the experiment. PEEP was set at 0 cmH_2_O. 2) <u>Hyperoxia</u>: FiO_2_ was set to 100% during instrumentation and continued for the duration of the experiment. 3) <u>Instillation of gastric particles into the lungs</u>: Prior to experiments, a uniform stock of gastric contents from healthy donor pigs was homogenized in sterile saline, strained, filtered, and autoclaved. A sufficient volume was made for the planned experiments and was stored (−20°C) until use. At the time of experimentation, the gastric particles were resuspended in saline (40 mg/mL) with a pH of 1 as previously described(25, 26). Six aliquots (8 mL each), were bronchoscopically-instilled to lobar bronchi 15 minutes following the sham renal inoculation. 4) <u>Sham renal inoculation</u>: A 5 mL aliquot of Phosphate Buffered Saline (PBS) was administered into the kidney parenchyma as described above.

#### Group 3 – *Combined indirect and direct lung injury*

Both direct and indirect insults were used in this group in the order of 1-2) volutrauma and hyperoxia, 3) *E. coli* renal inoculation, and 4) bronchoscopic instillation of acidified gastric particles.

### Monitoring and data collection

Animals were monitored for at least 12 hours after renal inoculation. Sequential hemodynamic measurements including MAP, PAP, HR, and temperature were monitored and recorded continuously (MP160, Biopac Inc. Goleta, CA). Ventilation parameters including peak airway pressure (AP_peak_), respiratory rate, FiO_2_, and PetCO_2_ were recorded every hour along with arterial and mixed venous blood samples. Additional blood and and chest radiographs were obtained every 4-6 hours. At the conclusion of the experiment, animals were euthanized while under general anesthesia using intravenous potassium chloride (1-2 meq/mL). Organ tissue samples were acquired for histological assessment by an expert Pathologist. The chest radiographs were scored by a blinded Pulmonary & Critical Care-trained physician (MWS), who rated each chest radiograph on a scale of 1-10 to quantify the extent of lung injury (1 = no abnormalities, 10 = severe, diffuse lung injury.

### Prespecified criteria for successful model development

Prior to initial experimentation, the study team agreed to prespecified criteria by which the model would be considered a successful model of human ARDS: 1) recapitulation of the physiologic and histopathologic features of human ARDS: impaired oxygenation (PaO_2_/FiO_2_ < 300) and pathologic evidence of diffuse alveolar damage; 2) a time-efficient model in which ARDS is achieved within 24 hours of initial exposure.

### Statistical analyses

Descriptive data are presented as means and standard deviation (SD). Analysis of variance with repeated measures (RM-ANOVA) or mixed-effects analysis (in case of missing data) were used for longitudinal analysis including a post-hoc Tukey correction for multiple comparisons. primary analysis was conducted at the 12 hour mark. For animals who reached the pre-specified stopping criteria and were euthanized prior to 12 hours, their last recorded value was used. All data and figures were analyzed and created using Matlab R2017a (The MathWorks, Inc., Natick, MA), SAS 9.4 (version 9.4, Cary, NC), and PRISM 8 (GraphPad Software, San Diego, CA).

## Results

Baseline characteristics of animals in all experimental groups are presented in **Table 1**. Of the 14 animals, 11 survived to 12 hours for all measurements while 3 met prespecified criteria prior to 12 hours. Two of these were in the *indirect lung injury* group (9 and 11 hours of measurement) and one was in the *combined lung injury* group (11 hours of measurements).

### Oxygenation

We first compared oxygenation over time across the experimental groups as assessed by PaO_2_/FiO_2_ ratio (5, 7). As shown in **Figure 1**, experimental groups had similar baseline oxygenation. However, over time, Groups 1 (*indirect lung injury only*) and 2 (*direct lung injury only*) exhibited mild impairment in oxygenation, with mean PaO_2_/FiO_2_ plateauing at or above the definitional threshold of 300. In contrast, Group 3 (*combined indirect and direct lung injury*) exhibited a progressive decline in PaO_2_/FiO_2_ ratio from 494.1(67.46) at baseline to 124.8(63.80) at hour 12 (*P*=0.0012). While within-group variation was observed, all animals in Group 3 (*combined indirect and direct lung injury*) reached the definitional PaO_2_/FiO_2_ ratio threshold of ≤ 300 by hour 12. We thus concluded that the *combined indirect and direct lung injury* exposures provoke a level of impaired oxygenation that is consistent with human ARDS(5).

**Figure 1.**
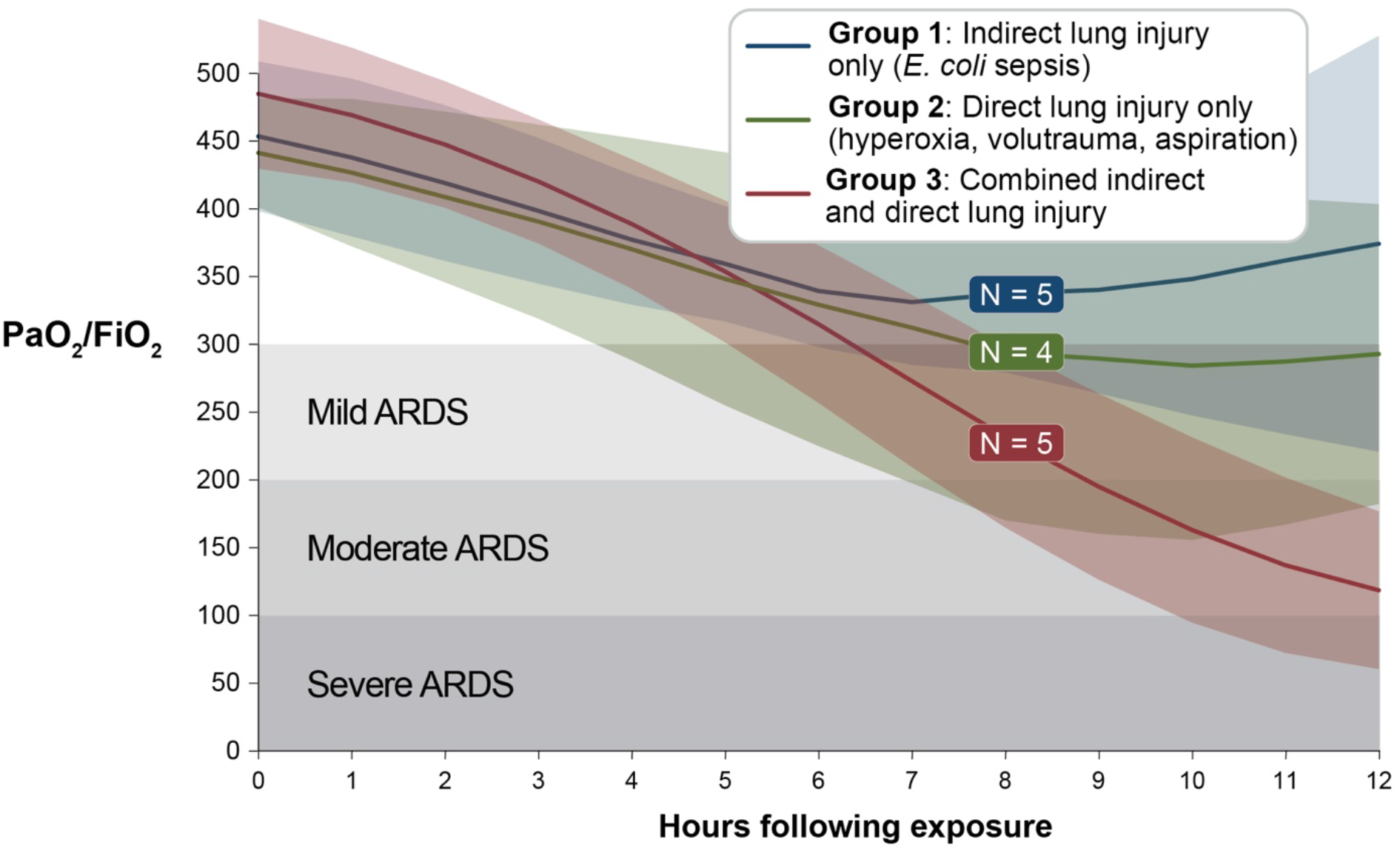
Comparison of oxygenation across experimental groups. Healthy Yorkshire-mix swine, 14-16 weeks of age, were exposed to 1) *indirect lung injury* (*E. coli* sepsis), 2) *direct lung injury* (hyperoxia, volutrauma, and aspiration of gastric particles), and 3) *combined direct and indirect lung injury* (all above exposures). Lines and variance represent means and standard deviation, both with Lowess smoothing.

### Chest imaging

We next compared serial chest radiographs from the animals in each experimental group (**Figure 2**). We specifically assessed for the presence of bilateral opacities, another definitional feature of ARDS(5). Chest radiographs were scored by a Pulmonary & Critical Care-trained physician, blinded to experimental group and timepoint, using a scale of 1-10 (1 = no abnormalities, 10 = severe diffuse bilateral opacities). Chest radiographs were obtained on a single animal in Group 1 (*indirect lung injury only*); these images were scored as normal (score = 1) both at baseline at hour 12. In contrast, both Group 2 (*direct lung injury only*) and 3 (*combined direct and indirect lung injury*) animals exhibited significant increases in chest radiograph abnormalities. In both groups, all baseline radiographs were scored as normal with a range of 1-2. In Group 2 (*direct lung injury only*), the chest radiograph score increased to a mean of 6 (SD 2.1) (range 3-9, 95% CI: 2.1, 9.9). Group 3 (*combined indirect and direct lung injury*) increased to a mean of 7.4 (SD 4.3 (range 5-10, 95% CI: 4.3, 10.5). As a test of internal validity, we compared the severity of impaired oxygen (PaO_2_/FiO_2_ ratio) and severity of injury on chest radiographs (chest radiograph severity score). Mixed effects regression confirmed that an increased chest radiograph severity score was significantly associated with decreased PaO_2_/FiO_2_ ratio when adjusted for experimental group and time point (*P*=0.008). These data demonstrate that the *combined indirect and direct lung injury* exposures result in the development of diffuse bilateral pulmonary infiltrates that are consistent with the human ARDS definition(5).

**Figure 2.**
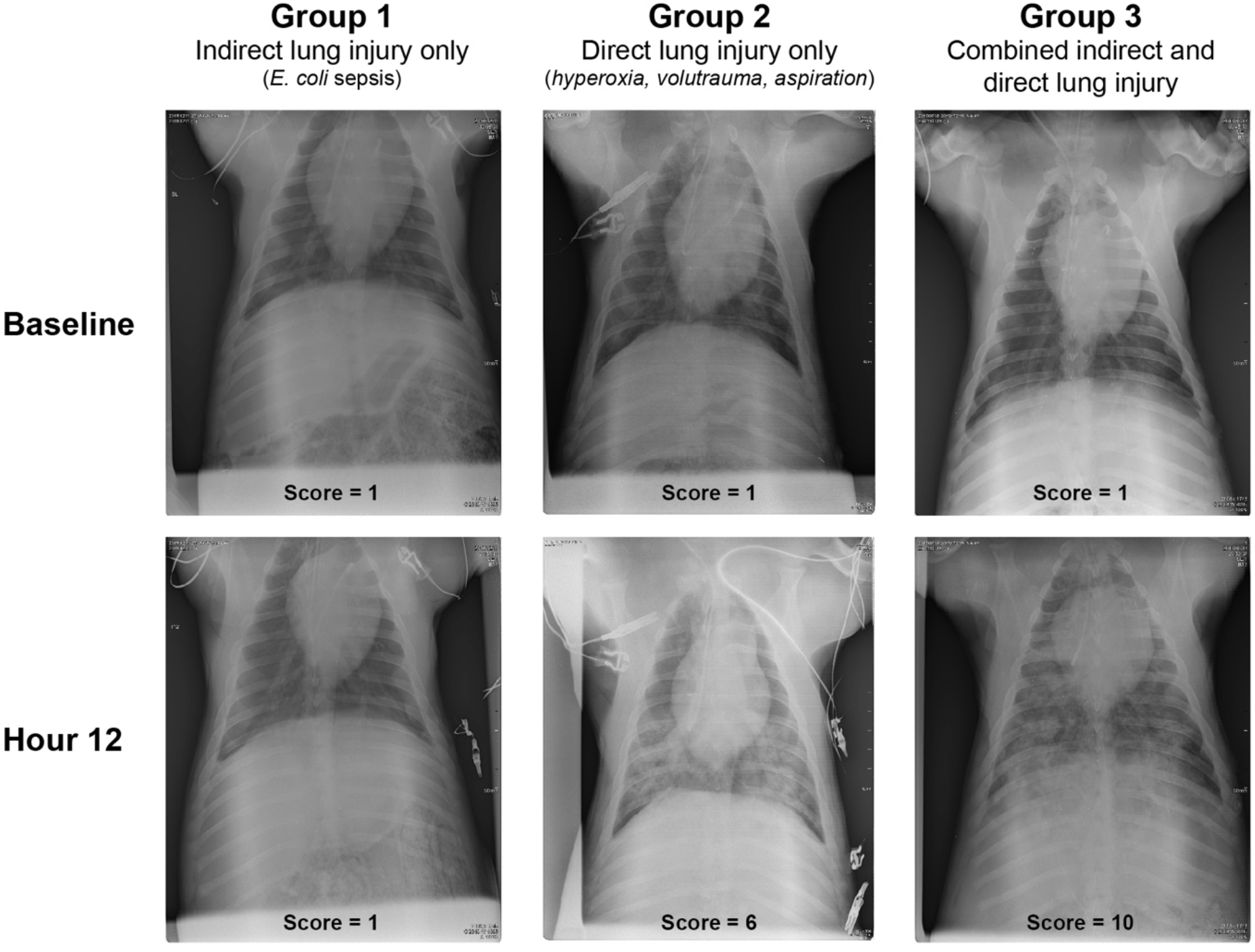
Representative chest radiographs across experimental groups. Ventral-dorsal chest radiographs were obtained at baseline and every 4 hours following exposure for the duration of the experiment. Images were scored by a Pulmonary & Critical Care Medicine physician (blinded to experimental group and timepoint) using a scale from 1 (normal) to 10 (severe, diffuse bilateral opacities).

### Histopathology

The histopathology of post-mortem lung tissue from the three experimental groups was assessed by an expert thoracic pathologist (**Figure 3**). None of the specimens from Group 1 (*indirect lung injury only,* n = 4) or Group 2 (*direct lung injury*, n = 4) exhibited the core features of DAD, including hyaline membrane formation (the histopathological hallmark of DAD). In contrast, in Group 3 (*combined indirect and direct lung injury*, n = 5), lung tissue from four of five animals met criteria for *definite DAD* based on the presence of hyaline membranes and other key features (e.g. intra-alveolar edema, fibrin thrombi). Within Group 1 (*indirect lung injury only*), three of four examined lungs were histologically graded as *normal*, with a single animal exhibiting increased interstitial cellularity and focal acute pneumonia. Within Group 2, four of four examined lungs were characterized by acute bronchopneumonia with intra-alveolar edema. As such, the *combined indirect and direct lung injury* exposures resulted in DAD, whereas the individual *indirect lung injury* and *direct lung injury* exposures did not.

**Figure 3.**
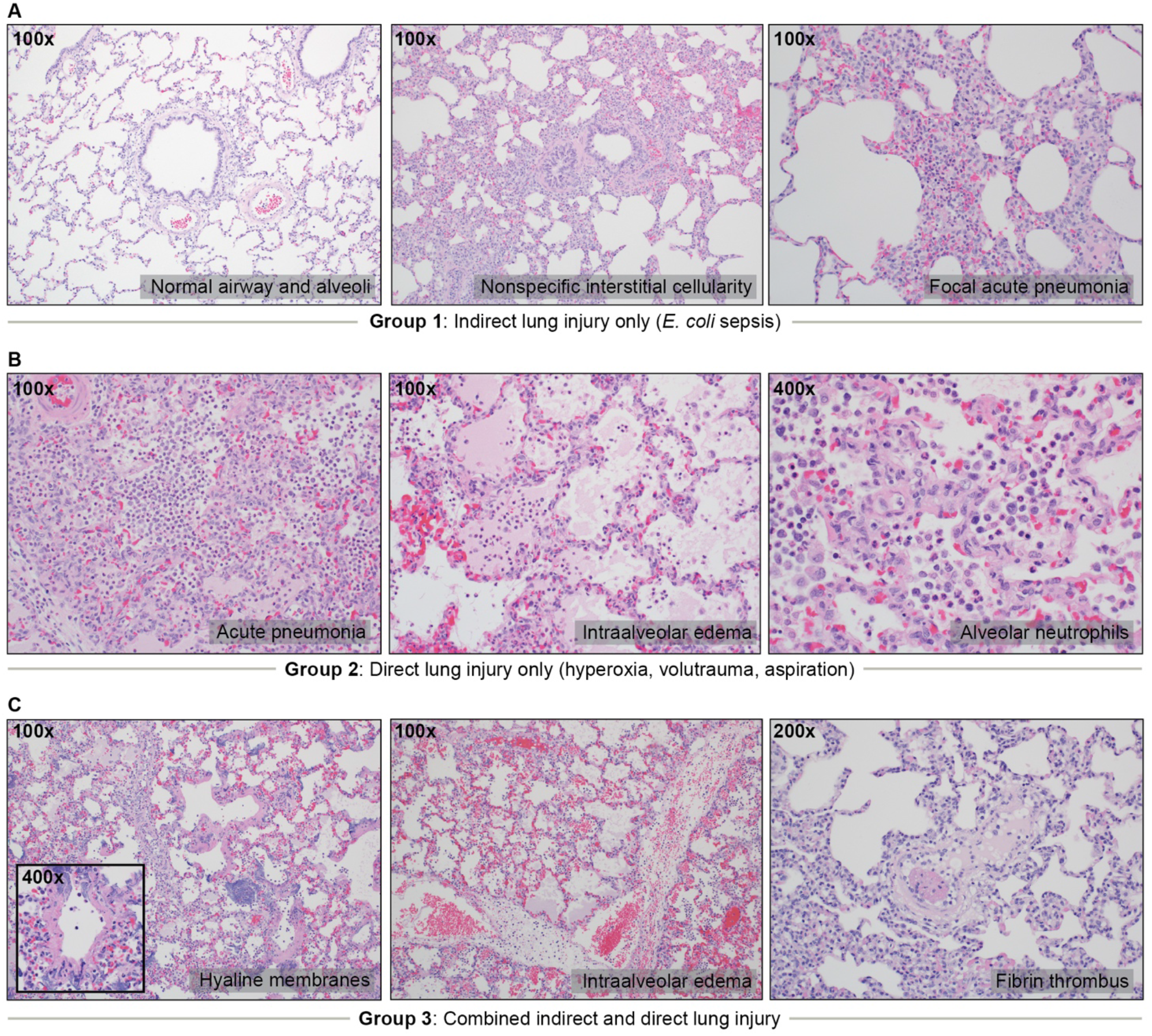
Representative histopathology across experimental groups. Post-mortem lung tissue was examined by an expert thoracic pathologist using a semi-quantitative instrument for identifying key features of Diffuse Alveolar Damage (DAD). *(A)* Of the four animals examined in Group 1 (indirect lung injury only), three were graded as *normal.* Abnormal findings included mildly increased interstitial cellularity and focal acute pneumonia in a single animal. No animals in Group 1 exhibited features of DAD. (*B*) Of the four animals examined in Group 2 (direct lung injury only), all four exhibited features of acute bronchopneumonia with intra-alveolar edema. No animals in Group 2 exhibited features of DAD. (*C*) Of the five animals examined in Group 3 (combined indirect and direct lung injury), 4/5 were classified as *definite DAD.* Prominent findings in Group 3 included hyaline membranes (4/5), intraalveolar edema (3/5), fibrin thrombi (5/5), and acute bronchopneumonia (5/5).

### Physiologic, Inflammatory, and Extrapulmonary Organ Function Measurements

Additional data regarding physiologic, immunologic, and extrapulmonary organ function measurements are included in the **Online Supplement**. At hour 12, peak airway pressures were increased in Group 2 (*direct lung injury only*) and Group 3 (*combined indirect and direct lung injury*) relative to Group 1 (*indirect lung injury only*) (**Supplemental Figure 1**). Arterial carbon dioxide (PaCO_2_) was greater in Group 3 (*combined indirect and direct lung injury*) than in Group 1 (*indirect lung injury only*). Experimental groups did not differ at hour 12 in their white blood cell count or relative neutrophilia (**Supplemental Figure 2**). Groups 1 (*indirect lung injury only*) and 3 (*combined indirect and direct lung injury*) both exhibited biochemical evidence of acute kidney injury (**Supplemental Figure 2**).

## Discussion

We here report a novel swine model of ARDS that faithfully recapitulates the features of human disease using common, clinically relevant injury exposures. Our model meets our prespecified criteria for successful model development for human ARDS: 1) it recapitulates the physiologic and histopathologic features of human disease (impaired oxygenation and diffuse alveolar damage) and 2) it does so in a time-efficient manner in which ARDS is achieved within 24 hours of exposure. Our novel model offers advantages over both small animal (rodent) models as well as existing swine models that rely on clinically unrepresentative single-hit exposures (e.g. oleic acid infusion(18, 19), surfactant washout(20)). Additionally, in alignment with NHLBI clinical research priorities, our novel preclinical model 1) uses a biologically-relevant infectious exposure (inoculation of viable *E. coli*)(23), 2) allows for the study of organ dysfunction and organ support, and 3) permits cointerventions (e.g. intravenous fluids, vasopressors, and antimicrobials). Our model thus fills an important gap in the preclinical study of ARDS, a devastating and common condition for which we lack molecular diagnostics and therapeutics.

In addition to meeting our own prespecified criteria, our model meets other established criteria for ARDS. Our model consistently provokes DAD (including hyaline membrane formation), the histopathological hallmark of human ARDS. This pathological finding is highly specific, and confirms that the model’s hypoxemia and radiographic opacities are not attributable to competing processes (e.g. shock, cardiogenic edema, or acute pneumonia). Our model also satisfies the clinically-derived Berlin Criteria(5), which are typically considered inapplicable to animal models given the impracticality of assessing arterial oxygenation and chest radiographs in rodents(7). Finally, our model fulfills criteria established by a 2011 American Thoracic Society workshop on experimental lung injury in animals(7), in that it induces 1) severe lung injury within 24 hours of exposure, 2) histologic evidence of tissue injury (e.g. hyaline membranes), 3) alteration of the alveolar capillary barrier (e.g. proteinaceous edema within the alveolar space), 4) alveolar inflammation (e.g. accumulation of alveolar neutrophils), and 5) physiologic dysfunction (e.g. hypoxemia). In aggregate, our model thus robustly satisfies pathological, clinical, and experimental criteria for ARDS.

Importantly, these criteria for modeling ARDS were only met by our *combined* exposure group (both *indirect lung injury* and *direct lung injury* exposures), and were not met by its individual constituent exposures (*indirect lung injury only* or *direct lung injury only*). These findings are congruent with recurring observations, both clinical(21) and experimental(22), that both epithelial *and* endothelial injury are necessary to yield the full pathophysiologic and clinical manifestations of ARDS. In the contemporary era, most patients with ARDS have risk factors that represent both systemic (endothelial) pathology (e.g. sepsis or shock) as well as direct lung injury (e.g. pneumonia or aspiration)(21). Even SARS-CoV-2, a pandemic respiratory virus that causes ARDS in its most severe form, provokes both epithelial *and* endothelial lung injury as assessed via post-mortem examination(27, 28). We believe this trend has important experimental implications, as “single-hit exposures” (such as intratracheal endotoxin in mice) are unlikely to fully recapitulate the complex pathophysiology of human ARDS. Strong consideration should be given to leveraging combined exposures to improve the biological and clinical relevance of experimental lung injury.

We acknowledge that our study has several limitations that should inform future investigations. Firstly, this pilot study used only male animals to minimize one source of biologic heterogeneity. Future studies will include both male and female animals to investigate the role of sex in the susceptibility to ARDS(29). Secondly, we did not include a negative control arm (i.e., animals that received either sham indirect and/or direct lung injury exposures). Despite this, our *indirect lung injury only* experimental group exhibited near-normal lung histopathology, providing reassurance that supportive care and instrumentation alone are not responsible for the ARDS pathophysiology observed in our combined exposure group. Additionally, the availability of serial sampling afforded by large animal modeling permitted us to perform within-group comparisons to baseline (pre-exposure) measurements for physiologic and radiographic features. Finally, we did not design the model to test differences in survival, long-term management, or sequelæ. In future studies, our model will serve as a foundation to test interventions such as supportive care (e.g. ventilation strategies) or pharmacotherapy.

In conclusion, we report a novel high-fidelity swine model of ARDS provoked by common, clinically-relevant injury exposures. As a controlled large animal model, it permits longitudinal measurement of physiologic, radiographic, and biochemical features of disease, as well as definitive histopathologic evaluation of lung tissue. This model fills an important pre-clinical gap in the study of ARDS, and will facilitate translational inquiry into the pathogenesis and management of this lethal and common lung condition.

## Acknowledgement

The authors would like to thank Christopher Altheim, Christopher Fry and the staff of Michigan Center for Integrative Research in Critical Care (MCIRCC) for their technical support.

---

**Supplemental Figure 1.**
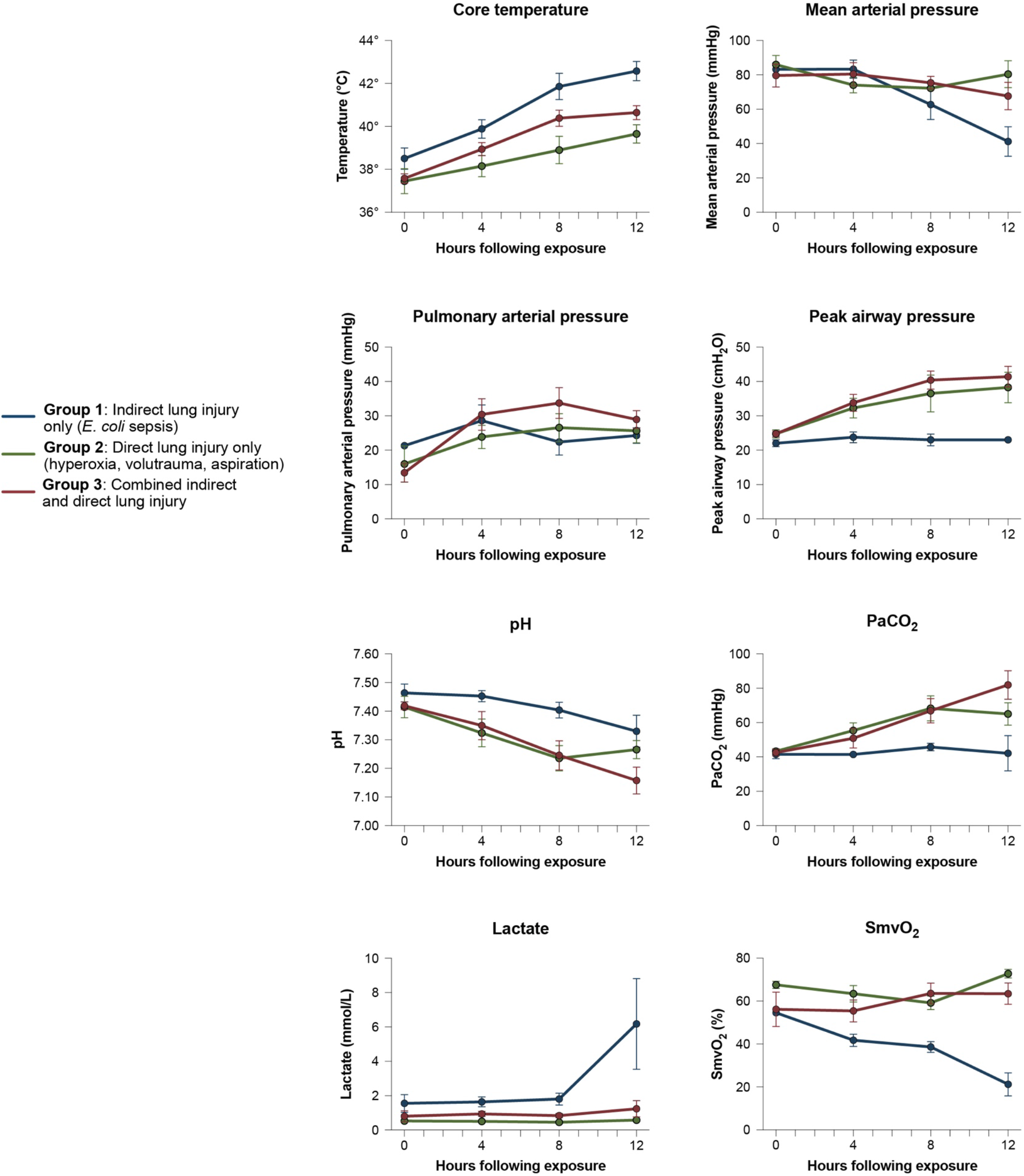
Comparison of physiology across experimental groups. Healthy Yorkshire-mix swine, 14-16 weeks of age, were exposed to 1) *indirect lung injury* (*E. coli* sepsis), 2) *direct lung injury* (hyperoxia, volutrauma, and aspiration of gastric particles), and 3) *combined direct and indirect lung injury* (all above exposures). Lines and variance represent means and standard deviation.

**Supplemental Figure 2.**
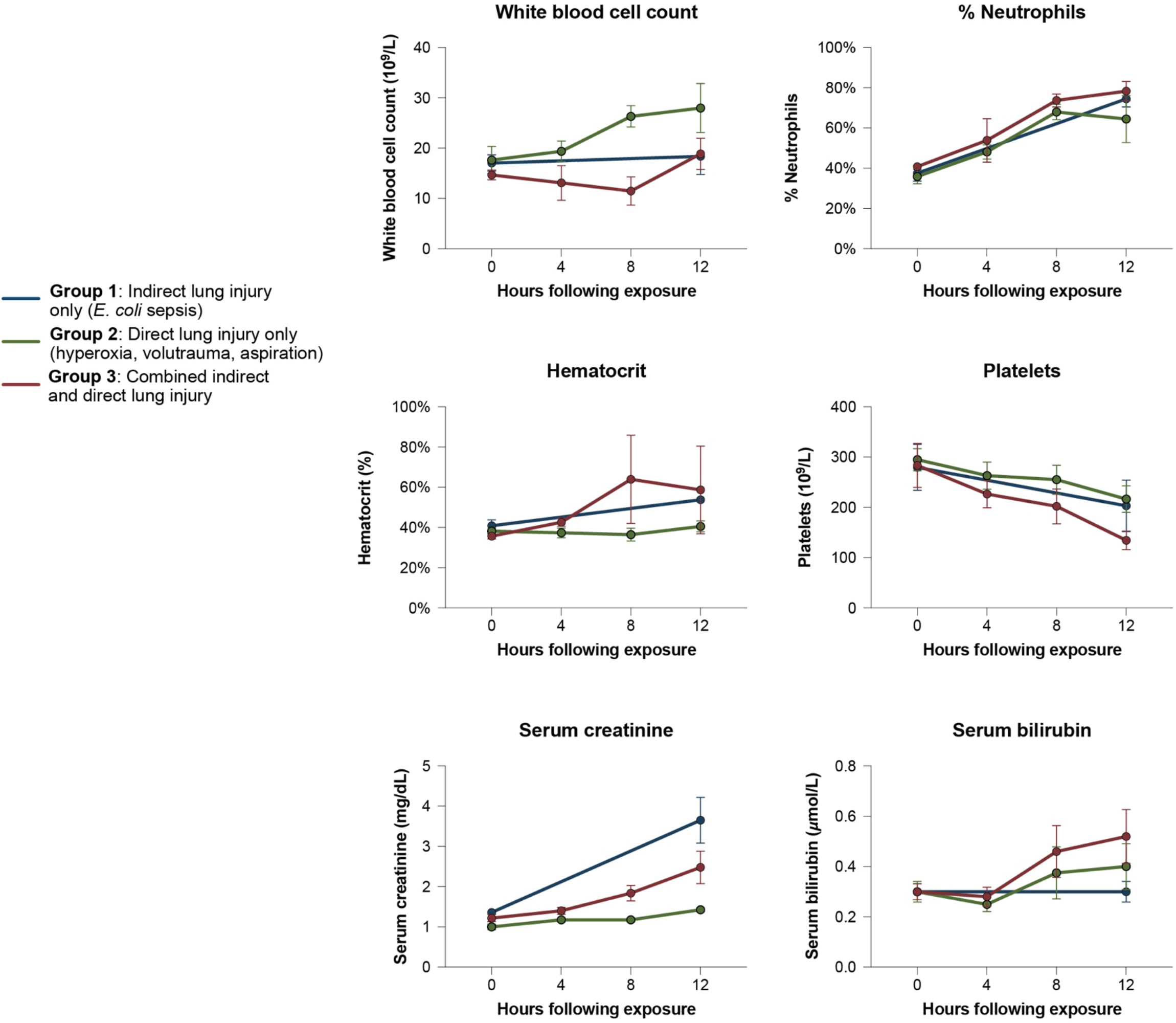
Comparison of laboratory values across experimental groups. Healthy Yorkshire-mix swine, 14-16 weeks of age, were exposed to 1) *indirect lung injury* (*E. coli* sepsis), 2) *direct lung injury* (hyperoxia, volutrauma, and aspiration of gastric particles), and 3) *combined direct and indirect lung injury* (all above exposures). Lines and variance represent means and standard deviation.

